# External structures at the nest site predict nest’s asymmetric architecture in mud-building birds

**DOI:** 10.1101/2025.08.14.670325

**Authors:** Nicolas M. Adreani, Victoria Morales Latorre, Lucia Mentesana

## Abstract

In the animal kingdom, nests are essential structures and textbook examples of extended phenotypes. However, the relationship between builders, nest traits, and the nest site remains poorly understood. We indirectly examine whether external components in the nest site influence nest building behavior, specifically focusing on their effect on nest architecture. We hypothesized that pre-existing structures at the nest site of horneros (Aves: *Furnarius rufus*) could influence nest construction, thereby affecting the nest’s architecture. To test this, we investigated the relationship between nest asymmetry—defined by the side on which the nest opening is located—and external lateral structures using a database of 12,356 nest photographs taken by citizen scientists across the bird’s entire distribution. We found that in nests built in contact with a lateral structure, birds were more likely to place the nest entrance on the same side as the lateral structure than by chance. When lateral structures were incorporated into the nest, this likelihood increased significantly. Our findings suggest that incorporating pre-existing elements into nests leads to predictable asymmetric architecture, highlighting a strong interaction between birds’ building behavior and nest site properties. These results underscore the importance of understanding the behavioral mechanisms shaping avian architecture and their plasticity.

## Introduction

The use of nests is widespread across insects, mammals, birds, fish and reptiles [1,2]. Varying in complexity, these animal structures are crucial as they provide a safe environment for early development and a shelter for adults [1,3]. Consequently, strong selection pressures are expected to act on both the form and function of nests, making them textbook examples of extended phenotypes [4,5]. Among the many animals that build nests, birds have emerged as key models for understanding the evolutionary, ecological, and adaptive significance of nest phenotypic traits [6–9]. In many species, cooperative nest-building between individuals can introduce additional social complexity to how these traits develop and evolve [10]. While the ecological and evolutionary drivers of avian nest phenotypic traits have become increasingly popular research topics [11,12], the potential effect that the nest site properties might have on the builders (or vice versa), and hence on the nest’s phenotypic traits, has received less attention.

The place where a bird chooses to build its nest is crucial. An incorrect assessment of nest site might lead to the selection of inadequate nest locations that can affect breeding success [13,14]. Structurally inadequate sites may also present challenges to the builders, both physically and cognitively, thereby influencing the nest’s architecture [7]. For example, house wrens (*Troglodytes aedon*) and blue tits (*Cyanistes caeruleus*) construct nests with traits that vary depending on the properties of the nest box, indicating that the interaction between builders and the environment affects nest architecture [15,16]. Research into this interaction has primarily focused on secondary cavity nesters (∼18% of bird species) using artificial nest boxes and often emphasizing nest materials [17,18]. For birds that must build their nests outside cavities, which account for most bird species, information on the interaction between the builders and the structural context of the nest during nest building is limited. For these species the physical properties of a potential nest site likely influence not only the birds’ nest site selection decision but also their building behavior, potentially affecting the nest’s final architecture, its stability and ultimately their reproductive success [19]. This raises several unresolved questions. For instance, if building behavior results from specific behavioral patterns, could the properties of the nest site constrain such patterns and, consequently, the nest’s architecture or the bird’s building capabilities? What is the degree of plasticity in birds that must adapt their building behavior to different nest site properties? Addressing these questions is difficult due to the need for standardized and comparable quantification of both nest site properties and nest phenotypic traits in the wild. Furthermore, many nest traits are labile and difficult to measure without damaging the nest, particularly in stick-based nests.

We identified a nest phenotypic trait that integrally describes the nest’s architecture and can be quantified unequivocally in the wild: bilateral asymmetry [20]. Male and female horneros (*Furnarius rufus*; Fig. 1A) cooperatively build a complex and conspicuous mud nest that is bilaterally asymmetric (Fig. 1B) [20–23]. When the nest opening is on the left (i.e., ‘left asymmetry’), the incubation chamber is located on the right side of the dome and the opposite when the nest entrance is located on the right (i.e., ‘right asymmetry). Bilateral asymmetry does not occur randomly in this species, does not seem to be explained by macro-ecological factors or micro-habitat properties, and is highly repeatable among pairs (repeatability = 0.65) [20]. This aspect of hornero’s nest architecture presents a clearly-defined binomial phenotypic trait that can be observed and quantified in the wild. When horneros build their nest, they can build it isolated (Fig. 1A) or in contact with a lateral structure, which can be a tree, an anthropogenic structure like a wall, or even another hornero nest from previous breeding seasons (Fig. 1C). Here, contact between the nest and the lateral structure can be tangential, or the lateral structure can replace parts of the nest, which can save materials, time and energy needed for construction (Fig. 1D). When contact is tangential, the birds interact with the lateral structure at the beginning of the building process, but if the pre-existing structure replaces a part of the nest, it unequivocally interferes with the normal building pattern that a bird would perform in the absence of such structure. Nest asymmetry in horneros is the outcome of a perceivably complex, yet inaccurately described, building process [24]. Thus, studying this nest trait in relation to properties of the nest site -like lateral structures- can be a reliable indirect indicator of whether external components of the nest site interfere or affect the nest building behavior of the birds.

**Figure 1.**
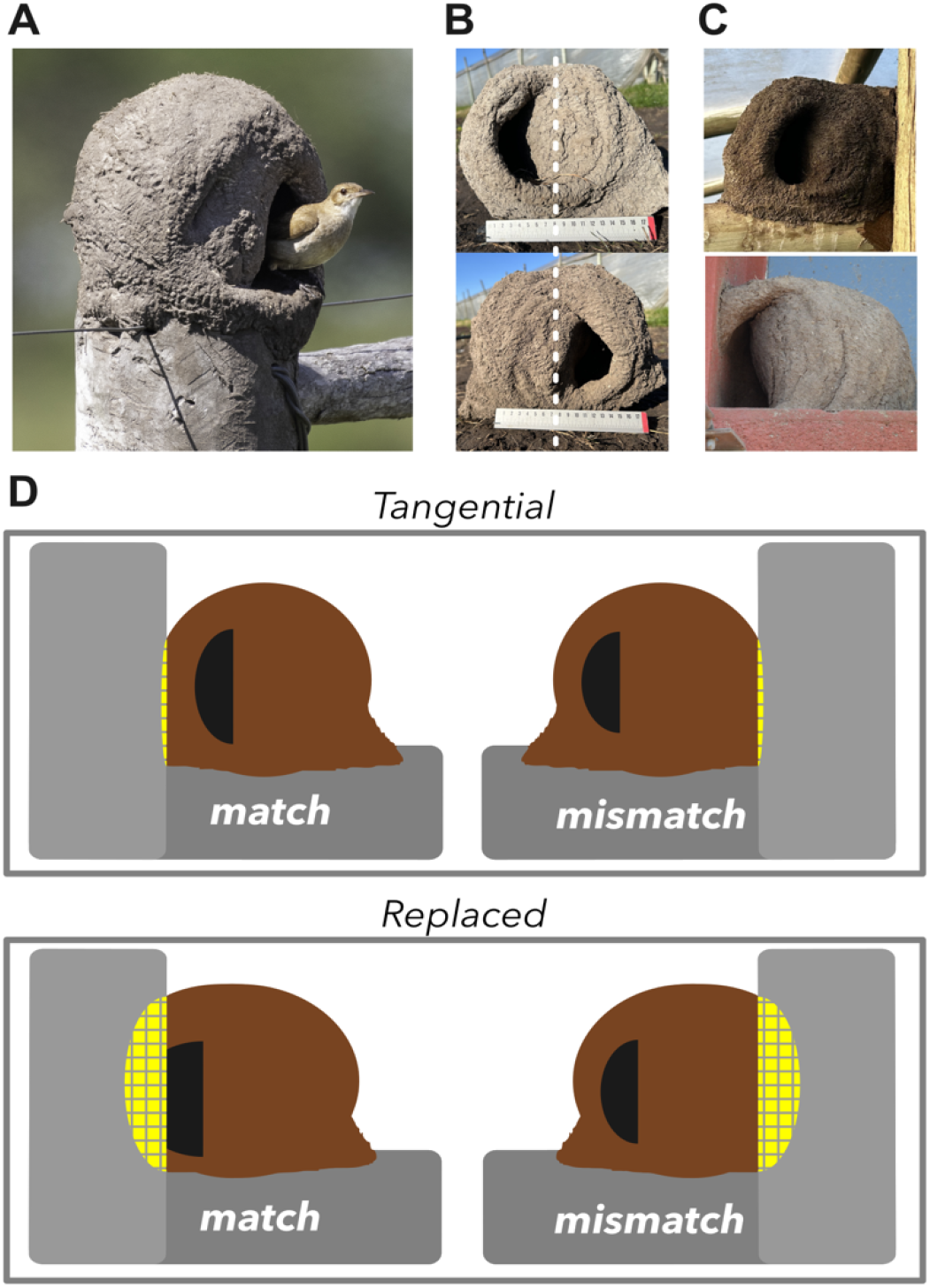
Nest entrance and lateral structures in hornero nests. **(A)** Hornero (*Furnarius rufus*) inside its nest. **(B)** Left and right asymmetric nests with a18 cm ruler for size reference and dotted vertical line showing bilateral asymmetry. **(C)** Nests in contact with a lateral structure. **(D)** Attachment can be ‘tangential’ when external nest walls are only in contact with the lateral structure, or ‘replaced’, when the lateral structure substitutes parts of the nest. In both cases there can be a match or a mismatch between asymmetry side and lateral structure side. When there is a match, both the nest asymmetry and the lateral structure are on the same side. When there is a mismatch, nest asymmetry and the lateral structure are on opposite sides. For clarity, we only show examples of nests with left asymmetry.

If horneros follow specific building rules that result in left or right asymmetry, we hypothesize that external structures at the nest site can interfere with such rules and, in consequence, influence the location of the nest entrance (i.e., left or right nest asymmetry). Thus, we predicted a relationship between the side of the external lateral structures at the nest site and the side of the nest opening. Our current understanding of hornero nest-building behavior does not allow us to accurately predict the relationship between the location of the nest entrance and the external lateral structures. Specifically, we cannot determine whether the nest entrance and lateral structures will be on the same side or opposite sides. However, any relationship between these elements should be independent of the side on which the lateral structure is located. Alternatively, if the interaction with external structures does not interfere with or influence the birds’ behavior, we predicted that the relationship between nest asymmetry and nest-site lateral structure should be like that expected by chance. When lateral structures are tangential to the nest, it is difficult to assess with precision the degree of interaction that the birds had with the structure during the building process. In contrast, when lateral structures replace part of the nest, we are certain that horneros must alter their building behavior by omitting parts of the nest wall. Therefore, we predict a stronger relationship between the side of the external structure and the side of the nest entrance when the pre-existing structures replace part of the nest compared to tangential cases. To test our predictions at the species level, we analyzed 12,365 pictures of hornero nests collected via a citizen-science initiative using a smartphone application. The data covered five countries where the species occurs (Argentina, Uruguay, Bolivia, Brazil, and Paraguay, Fig. 2A), spanning approximately 4.8 million km^2^—almost the entire species’ distribution.

**Figure 2.**
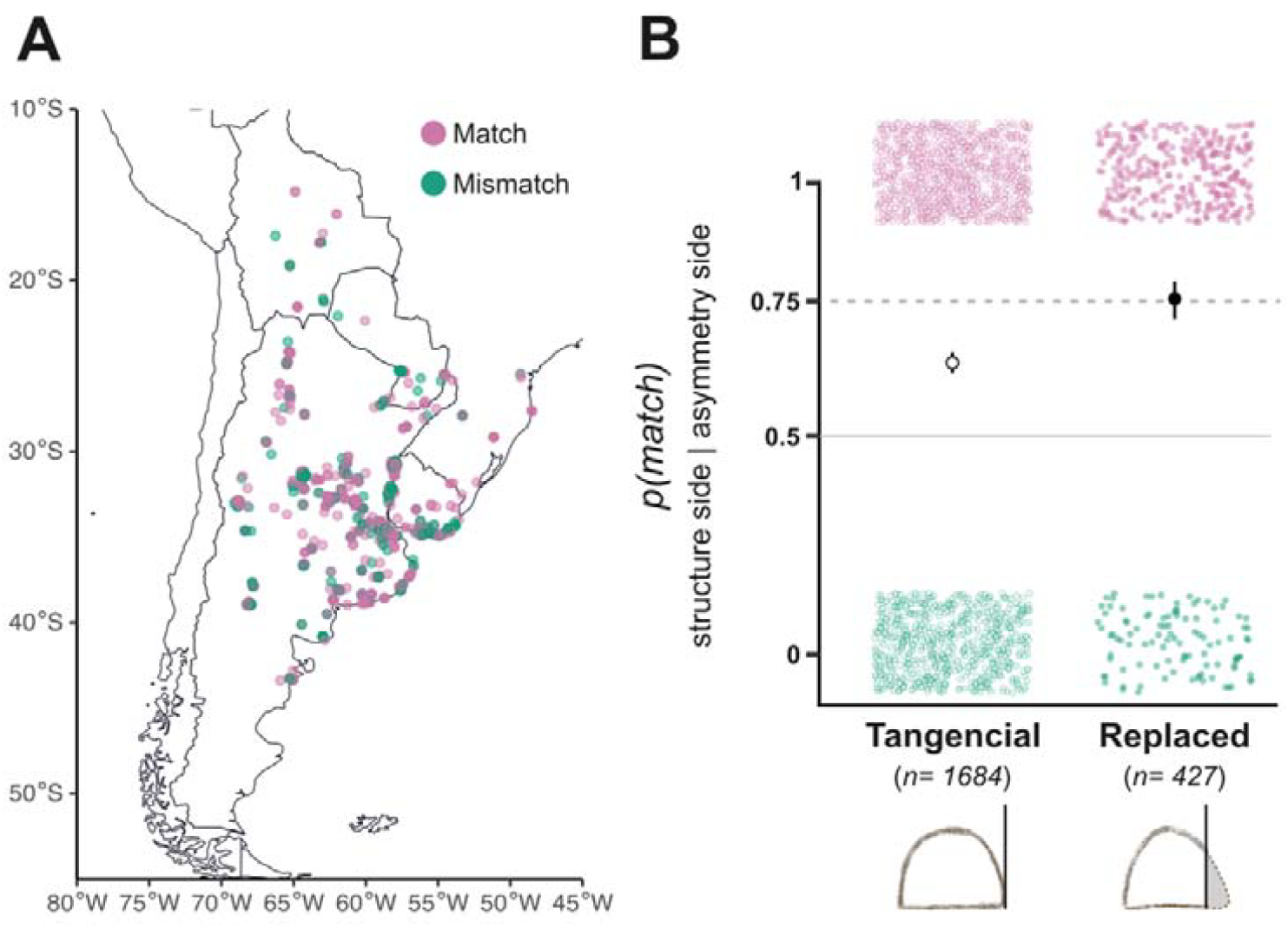
Relationship between lateral structures and location of the nest opening. **(A)** Spatial distribution of match (pink) vs. mismatch (green) cases. **(B)** Probability that the nest opening was on the same side as the lateral structure (*p(match)*) when the structure was tangential to the nest and when the structure replaced part of the nest. *p(match)*=1 refers to those cases where the lateral structure and the nest opening are on the same side and *p(match)*=0 to those where the lateral structure and the nest opening are on opposite sites. Point clouds are the raw data, black and white dots the back-transformed estimates from the binomial model (Table 1), and vertical bars the 95% credible intervals (CrI). The horizontal line at 0.5 represents the *p(match)* expected by chance. A statistical difference between estimates can be assumed when model estimates do not overlap the 95% CrI.

## Materials and Methods

### Data collection

We screened 12,365 photographs of hornero nests collected by citizen scientists *via* a free smartphone-application during one calendar year (October 2018-October 2019) [20]. Briefly, each user uploaded the picture of a nest *in situ* and noted different aspects of it, like nest asymmetry (left or right) and other micro-habitat features of the nest site (e.g., nest height, nest entrance orientation, urbanization context). In addition, cellphones with GPS sensors automatically registered the geolocation of each nest. In a citizen-science approach, pseudo-replication can be a concern, as multiple users might report the same nest. However, after conducting pair-wise distance comparisons between all nests, only 3.8% of them were found to be closer than 25 meters, the maximum resolution of the GPS sensors and a smaller spatial scale compared to horneros average territory size [20,23]. Therefore, if pseudo-replication occurs, it is minimal, especially because for this small proportion of nests it is more likely that other nests from the same pair were present in the territory than different individuals taking pictures of the same nest. For more details on data collection see [20].

**Table 1.** Results of the generalized linear model predicting the probability that the nest opening was on the same side as the lateral structure (i.e., *p(match)*). Explanatory variable was *Replacement* (Yes or No). We present fixed (β) parameters with their 95% credible intervals (CrI) in brackets.

|  |  |
| --- | --- |
| | $p$ (*match*) |
| *Fixed effects $\beta$*<br>(95% *CrI*) |  |
| Intercept (*Yes*) | 0.471<br>(0.374; 0.571) |
| No | 0.660<br>(0.414; 0.898) |

From all the pictures, we selected those nests that complied with the following criteria: nests were finished, the architectural asymmetry was clearly visible, and one of the nests’ lateral walls appeared to be in contact with a lateral structure (n = 2,782; Fig. 1C). We did not consider nests where laterals structures were present on both sides and we also excluded nests built adjacent to other nests, as in such cases, it was challenging to determine which nest was older or newer, and thus difficult to ascertain which served as an external structure to the other (n = 119 cases). Each nest was then assigned a reliability score from 0 to 2 concerning the contact of the nest with the lateral structure: ‘0’ difficult to assess if the nest was in contact with the lateral structure (n = 87); ‘1’ contact was highly likely but the extent of the contact could not be determined (n = 465); and ‘2’ the view of the lateral contact was clear and reliable (n = 2,111). To ensure data reliability, we only considered nests that had a reliability score of 2. For each of these nests we noted: 1) nest ID number; 2) asymmetry type (left or right); 3) side of the nest in contact with the external structure (left or right); and 4) whether the external lateral structure was tangential or was replacing parts of the nest wall (Fig. 1D). To identify potential geographical biasses of these nests, we visually explored the geographic distribution of 1916 nests with GPS location (Fig. 2A; we could not check the total 2011 pictures because some users did not have cellphones with GPS or the Location Services activated). In our previous study we found a small, yet statistically meaningful, spatial autocorrelation of nest asymmetry (i.e., for a small spatial scale of 0 to 2 km, nests with same asymmetry showed some degree of spatial clustering). We do not expect spatial autocorrelation to be a confounding factor in this study, as nests with lateral structures are widely distributed and not spatially clustered in a 0 to 2 km scale (Fig. 2A).

### Statistical analyses

Analyses were performed in R v.4.3.2 (R Core Team 2015) with the packages ‘lme4’ [25] and ‘arm’ [26] using a pseudo-Bayesian framework with non-informative priors [27]. Linear model assumptions were verified by inspecting residual plots using the package ‘DHARMa’ [28]. We used the ‘sim’ function to simulate posterior distributions, extracting mean estimates, 95% credible intervals, and posterior probabilities from 10,000 simulations [29]. Effects were considered statistically meaningful when zero was not included in the 95% credible intervals [27].

To test our hypothesis, we ran one generalized linear model with binomial distribution. We created a binomial response variable ‘*p(match)*’ that was defined as 1 if the side of the external structure and the side of the nest entrance were the same (i.e., ‘match’) and 0 if the two were at opposite sides (i.e., a ‘mismatch’; Fig. 1D). We considered one categorical explanatory variable: i) ‘*replacement*’, defined as ‘no’ if the structure would only be tangential to the nest walls, and ‘yes’, if the pre-existing structure was replacing part of the nest walls (Fig. 1D). **Results**

From the 12,365 nests in our database, 17.0% (n = 2,111) were in contact with one lateral structure. From this subset, in 20.1% (n = 427) the lateral structure replaced part of the nest walls.

When the nests were in tangential contact with a lateral structure, the probability that the side of the nest entrance was on the same side as the structure (*p(match)=1*) was 0.62, significantly larger than the 0.50 probability expected by chance (Fig. 2B; Table 1). Here, there was a marginal statistical difference in this effect comparing left and right asymmetry nests (Fig. S1; Table S1).

When the pre-existing structure was replacing a part of the nest, the probability that the side of the nest entrance was on the same side as the structure was of 0.76, also significantly larger than the 0.50 probability expected by chance and 22.1% higher than in tangential cases (i.e., comparing the probability of 0.62 in tangential cases vs. that of 0.76 in replaced cases; Fig. 2B; Table 1). Here, there was no statistical difference in the effect comparing left and right asymmetry nests (Fig. S1; Table S1).

## Discussion

We found an association between the side of pre-existing lateral structures and the side at which the nest opening was located. When the nest was tangentially in contact with a lateral structure, the probability of the nest opening being on the same side of the structure was 21% higher than expected by chance (Fig. 2B). When the pre- existing structure replaced a part of the birds’ nest, the probability of the nest opening being on the same side of the structure was 42% higher than expected by chance (Fig. 2B). This pattern implies that, on average, when building their nests three out of four pairs of horneros avoid contact between the lateral structure and the incubation chamber when the external structure replaces part of the nest.

Building a nest in contact with lateral structures may offer several advantages. First, lateral structures could increase the nest’s stability against harsh weather, which can destroy nests and cause breeding failure [30,31]. Second, lateral structures might also enhance nest concealment, potentially reducing predation or brood parasitism [32,33]. Third, incorporating pre-existing structures might also benefit the parents if it reduces the energy and time required for nest construction [8,34]. Despite these potential benefits, only 2,111 of the 12,365 nests examined (17.2%) were attached to lateral structures. Of those, only 427 (3.45% of the total nests) incorporated pre- existing structures into their nest. These low proportions suggest that attaching the nest to a lateral structure is not a common strategy and, when chosen, incorporating the laterals structures does not seem to be a preferred option either. With so many potential advantages, why are these choices uncommon?

From a behavioral standpoint, it is possible that the external structure interferes with the bird’s behavior and only a small proportion of hornero pairs can cope with the external structure. When there is a lateral structure incorporated into the nest, the birds must -inevitably- interact physically with the structure and adapt the way they build their nest in comparison to when no structure is present. This is something that many builders, including humans, encounter and requires behavioral plasticity. For example, several species of ants adapt their nest traits to properties of the nest site [35]. Hornero pairs tend to build nests with the same architectural asymmetry in consecutive nest building events, suggesting either a hard-wired genetic basis or the consequence of learning of such nest phenotypic trait [20]. The strong relationship between nest asymmetry and pre-existing structures that we uncover indicates that the lateral structure must influence the bird’s behavior in some way. Horneros might be able to build a nest with one or the other asymmetric architecture only, or birds might have a predisposition or preference towards building a nest with a certain asymmetry, something that could be compared to human or animal handedness [36,37]. In either case, it is possible that horneros follow specific nest building rules that are tightly linked with the nest’s final asymmetric architecture. Here, the presence of an external structure might interfere with such behavioral pattern in a way that the most efficient, most comfortable or simply the compatible way to build the nest incorporating a lateral structure is if the nest opening is placed on the same side of the lateral structure. Despite the relationship between the nest asymmetry side and the lateral structure is strong, in ca. 25% of the cases hornero pairs seemed capable of building the nest opening opposite to this pattern. This could be explained: i) by some birds being more plastic; ii) by the variation in the degree of interaction between the builders and the lateral structures in our data or iii) by a compensatory mechanism related to the cooperative aspect of nest building (i.e., within hornero pairs, one member of the pair might have a compatible building predisposition that fits with the position of the external structure). Yet, several questions emerge. First, are the birds actively choosing to incorporate the structure into their nest, or is this a sign of plasticity in adapting to the presence of a lateral structure, which could also become a limitation? Second, are some birds more plastic than others in response to the presence of the lateral structure and can build their nest in any way? Third, do birds choose or prefer sites with a laterals structure placed such that it fits their preferred asymmetry type?

From an ecological standpoint other possibilities emerge, related to the cooperative aspect of hornero’s nest building. Pair compatibility has been suggested as an important success factor within socially monogamous birds (e.g., [38]). In horneros a successful nest building event might depend on properties that emerge from the interactions between partners. Here male and female horneros might -even differentially- be affected by the presence of the lateral structure. In biparental bird species, cooperation and synchrony, can lead to increased reproductive success and can even be important for coordinating male and female’s reproductive physiology (e.g., [39]). In this context, importance of nest building and nesting displays has been discussed, and higher success is expected in better coordinated pairs [40]. Hornero pairs might need to perform certain behaviors, or displays, during nest building that are important later for reproduction and that might be interrupted or difficulted by the presence of the lateral structures.

Incorporating an external structure into the nest may result from the birds’ active choice. Under this scenario, our results suggest that the birds would be expected to only choose a given nest location if there is a match between their asymmetry preference (or predisposition) and where the external structure is located. Another possibility is that the location of the nest at a site with an external structure is not a choice but rather a “miscalculation” or a ‘last option’ if no other site is available. Under these circumstances, the presence of an external structure could result in a limiting factor to building the nest if there is no match between the bird’s building capabilities (i.e., asymmetry preference or predisposition) and the location of the external structure. Both scenarios are consistent with the strong relationship between nest asymmetry and the pre-existing external structure that we find. Moreover, if birds actively choose to incorporate the structure into their nest, this would provide evidence of prospective cognition in hornero nest-building behavior [41], as the birds must anticipate the structure’s presence in their nest when choosing a site before or during construction. Whether such anticipation exists requires rigorous testing.

## Conclusion

Our study provides evidence that the interaction between the builder and the building site can have strong and measurable consequences for nest architecture. This is significant in the context of nest building processes, as our study reflects a type of ‘natural experiment’ where we indirectly evaluate how birds respond to the presence of a pre-existing structure during nest building by quantifying variation in asymmetric architecture. Our findings suggest there may be a behavioral or cognitive mechanism that prevents most birds from building their nests in such a way that the asymmetry is independent of the structural features of the nest site. Here, the cooperative nature of nest building in hornero breeding pairs adds complexity to understanding such interaction. Investigating the nest building process in detail is necessary to uncover possible behavioral rules underlying the construction, the role of cooperation, the possible role of lateralization, and how these interact to result in a nest. Unravelling the possible causes and mechanisms underlying the strong pattern we uncover represents an exciting avenue for novel empirical research in animal construction and architecture.

## Data availability statement

All the data and code to obtain these results are available on GitHub (https://github.com/mnadreani/article_asymmetry-nestsite). Upon acceptance of this work, the repository will be assigned with a DOI via Zenodo.

## Authors contribution

Conceptualization: NMA and LM. Methodology: NMA, VM and LM. Software: NMA. Validation: VM and MNA. Formal Analysis: NMA. Investigation: LM and NMA. Resources: LM and NMA. Data curation: MNA and VM. Writing – original draft: NMA. Writing – review & editing: all authors. Visualization: NMA and VM. Supervision: NMA and LM. Project administration: NMA and LM. Funding acquisition: NMA and LM.

## Notes

https://github.com/mnadreani/article_asymmetry-nestsite

https://data.mendeley.com/datasets/9745v8tj9h/1

### Competing Interest Statement

The authors have declared no competing interest.

